# TDFragMapper: a visualization tool for evaluating experimental parameters in top-down proteomics

**DOI:** 10.1101/2021.06.03.446879

**Authors:** Jonathan Dhenin, Diogo B Lima, Mathieu Dupré, Julia Chamot-Rooke

**Affiliations:** Mass Spectrometry for Biology Unit, CNRS, USR2000, Institut Pasteur, Paris, France; Université de Paris, Sorbonne Paris Cité, Paris, France; Leibniz-Forschungsinstitut für Molekulare Pharmakologie, Berlin, Germany

**Author notes:** The authors wish it to be known that, in their opinion, the first two authors should be regarded as joint First Authors.

## Abstract

**Motivation:** We present a new software-tool allowing an easy visualization of fragment ions and thus a rapid evaluation of key experimental parameters on the sequence coverage obtained for the MS/MS analysis of intact proteins. Our tool can deal with multiple fragmentation methods.

**Results:** We demonstrate that TDFragMapper can rapidly highlight the experimental fragmentation parameters that are critical to the characterization of intact proteins of various size using top-down proteomics.

**Availability:** TDFragMapper, a demonstration video and user tutorial are freely available at https://msbio.pas-teur.fr/tdfragmapper, for academic use; all data are thus available from the ProteomeXchange consortium (identifier PXD024643).

## 1 Introduction

Top-down proteomics (TDP) is a powerful technology allowing the characterization of proteins at the proteoform level using high-resolution tandem mass spectrometry (MS/MS). Proteoforms correspond to the different forms of a protein arising from all combinatorial sources of variation from a single gene (including combinations of genetic variation, alternative splicing, and post-translational modifications)(Smith and Kelleher, 2018). The complete characterization of proteoforms often requires the use of several complementary fragmentation techniques, such as collision-induced dissociation (CID), electron transfer dissociation (ETD) or ultraviolet photodissociation (UVPD)(Fornelli *et al*., 2018). In contrast to bottom-up proteomics, the experimental parameters used for the fragmentation in TDP, such as the activation energy or the charge state of the precursor ion chosen for fragmenting an intact protein, can significantly affect the quality of MS/MS data and therefore the protein sequence coverage (Brunner *et al*., 2015). Although existing tools are capable of matching a list of fragment ions to a protein sequence (ProSight Lite for instance)(Fellers *et al*., 2015), there is currently no computational tool allowing to visualize fragments arising from diverse MS/MS experiments on a unique fragmentation map without losing information on the contribution and the specificity of each experiment. Current tools provide a unique fragmentation map per MS/MS experiment, and thus their comparison, in particular when multiple parameters are assessed, is both difficult and time-consuming. Moreover, although the intensity of fragment ions can represent a precious source of information when dealing with MS/MS data, it is often not considered when matching fragments onto a protein sequence. To circumvent these limitations, we introduce TDFragMapper, a novel software-tool that can display and combine pre-assigned fragment ions achieved from various MS/MS experiments on a unique protein sequence, keeping an easy access to the individual contribution of each experiment and to the intensity of deconvoluted fragment ions. Our tool makes it possible to rapidly compare experimental parameters such as the type of fragmentation, the activation level or the precursor charge state in the MS/MS analysis of intact proteins. In what follows, we use TDFragMapper to evaluate the influence of essential experimental parameters in the TDP analysis of a standard mixture of intact proteins.

## 2 Material and Methods

To evaluate our software, we analyzed a standard mixture of six proteins (Thermo Scientific Pierce Intact Protein Standard Mix) in LC-MS/MS on an Orbitrap Fusion™ Lumos™ mass spectrometer using multiple experimental conditions, as described in the Supplementary Material. Raw files were deconvoluted using FreeStyle™ (v. 1.6.75.20) and the lists of deconvoluted ion masses were imported into ProSight Lite (v. 1.4) in order to assign fragments. Lists of assigned fragments were then exported and used with deconvoluted data as well as the protein sequence as input for TDFragMapper. The software was programmed in C# with .NET Framework 4.8 and requires a computer with Windows 10 or later, and at least 8 GB of RAM.

The objective of TDFragMapper is to propose an easy way to evaluate the individual contributions of key-parameters in TDP experiments in order to assess rapidly the combination leading to the best sequence coverage for intact proteins. The software is organized in three major interfaces: an input interface, a filter interface and a graphical visualization module.

Besides uploading data files, the input interface is used by the user to specify the putative protein sequence as a ^*^.fasta file and the experimental parameters used for each fragmentation experiment in an easy-to-fill table format. A nomenclature is proposed to ensure a standardized filling of these experimental parameters and an example of input format can be found by clicking on the help button. The drop-down menus and the arrow buttons of the filter interface allow the user to quickly set some experimental parameters and study their influence one by one (Figure 1, upper panel). Once filtered, the fragments are displayed in a visualization module and mapped onto the protein sequence (Figure 1, lower panel). Fragments are represented by vertical bars arranged in rows and colored according to the MS/MS experiment from which the fragment arises. N-terminal fragments (a-, b- and c-ions) and C-terminal fragments (x-, y- and z-ions) are displayed respectively above and under the protein sequence. A legend table is associated to each fragmentation map with the color code of fragments, the percentage of residue cleavages and the number of matching fragments for each experimental parameter under study.

**Figure 1:**
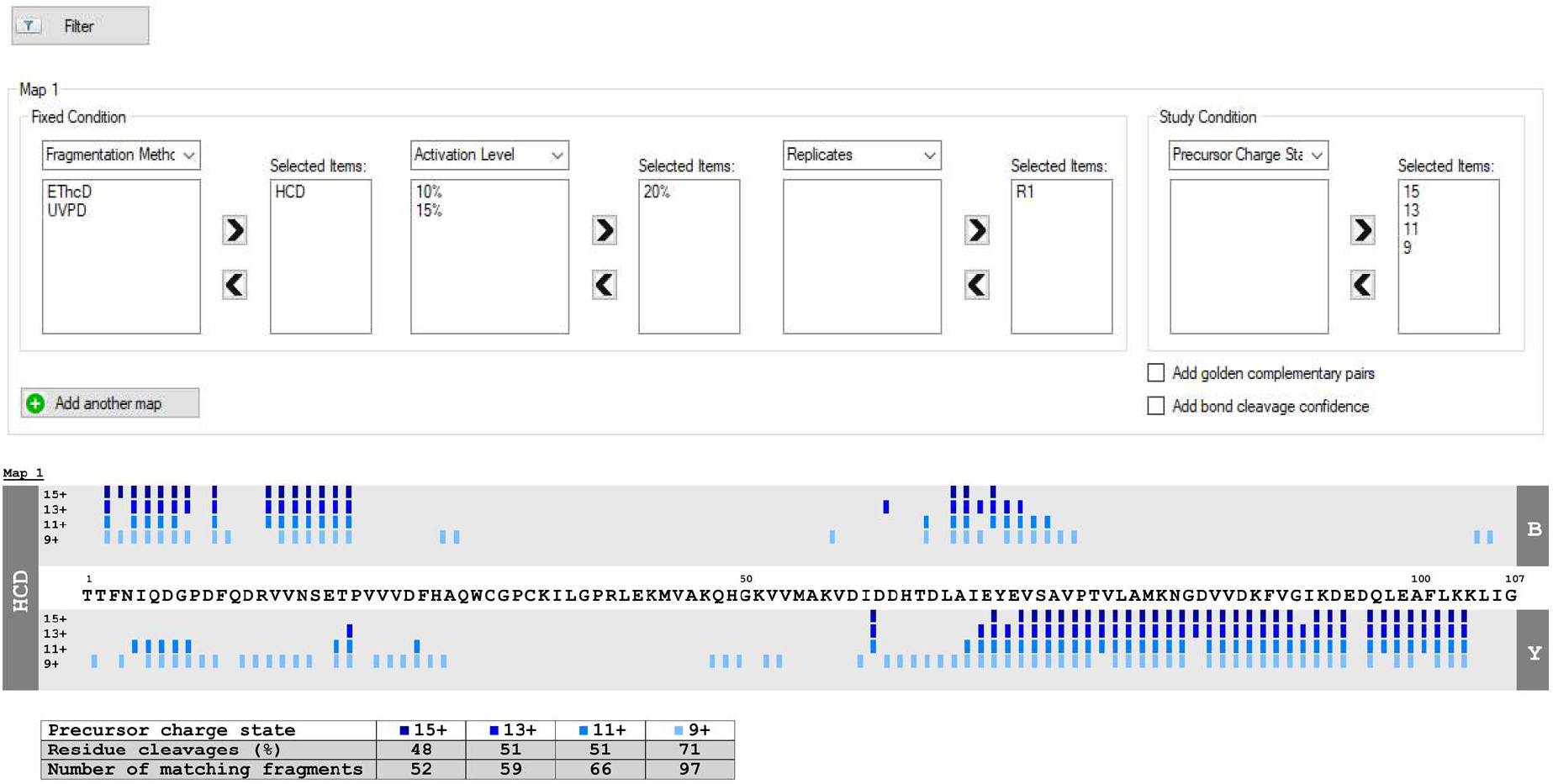
Screenshot of the filter interface of TDFragMapper (top panel) and the visualization module with the corresponding fragmentation map of human Thioredoxin obtained when varying the precursor charge state in HCD with NCE 20% (bottom panel)

## 3 Results & Discussion

Displaying fragments obtained from different MS/MS experiments onto a unique linear protein sequence with a specific color code makes it possible to highlight at a glance the contribution of each experimental parameter. TDFragMapper can be used for example to spot the dependency of a protein sequence coverage on the selected precursor charge state when using a particular fragmentation method (Figure 1, lower panel) or to assess the reproducibility of a particular fragmentation method across technical replicates. For instance, in Figure 1, it is easy to visualize that the charge state leading to the best sequence coverage for the HCD fragmentation of thioredoxin is the +9 one. Moreover, the user can add extra layer of information to the fragmentation map, such as the position and the number of golden complementary pairs (Kelleher *et al*., 1999; Horn *et al*., 2000)(Figures S1 and S2) or the intensity of deconvoluted fragment ions. This intensity option is easily accessible from the graphical visualization module and can be used to compare the abundance of common fragments across multiple MS/MS experiments on a single map (Figure S3). Once identified by the user thanks to the different fragmentation maps and the information provided by the previously described features, the results of the best fragmentation experiments can be summed using the merging option onto a single fragmentation map. This final map displays only the N-terminal and C-terminal cleavages regardless of the ion type and a final residue cleavage is computed, as can be seen in Figure S4.

Fragment maps can be exported as ^*^.tiff, ^*^.png or ^*^.jpg image files. TDFragMapper also allows to export a summary report as PDF^®^ file containing the information of uploaded data files and all the parameters used to create the maps. Finally, the work session and the results can be directly saved in an owner format (^*^.tdfm) that can be retrieved and used later. As far as we know, TDFragMapper has no equivalent and is highly complementary to other existing software tools allowing the analysis of targeted TDP datasets.

## 4 Final Remarks

We anticipate that TDFragMapper will ease the selection of optimal fragmentation parameters in order to increase the confidence in proteoform characterization in TDP experiments, including the precise localization of post-translational modifications. To facilitate its use, a tutorial with all functionalities is included in the tool’s website (Borges Lima *et al*., 2021) and can also be accessed through the help menu.

