## Supplementary Materials for "TDFragMapper: a visualization tool for evaluating experimental parameters in top-down proteomics"

* To whom correspondence should be addressed.

### Materials & Methods

#### Sample preparation

The Thermo Scientific™ Pierce™ Intact Protein standard mix is composed of the six following intact proteins: human IGF-I LR3 (9,111.47 Da), human Thioredoxin (11,865.52 Da), *Streptococcus dysgalactiae* Protein G (21,442.61 Da), Bovine Carbonic Anhydrase II (28,981.29), *Streptococcus* Protein AG (50,459.74 Da) and *Escherichia coli* Exo Klenow (68,001.15 Da). 76 µg of lyophilized protein mixture were reconstituted in 380 µL of solvent A (98% H2O, 2% ACN, 0.1% FA) to a final concentration of 0.2 µg/µL.

### LC-MS/MS

A Dionex UltiMate 3000 RSLC Nano System coupled to an Orbitrap Fusion™ Lumos™ mass spectrometer fitted with a nano-electrospray ionization source (Thermo-Scientific) was used. Five µL of protein sample were loaded at a flow rate of 10 µL.min$^{-1}$ onto an in-house packed $C_4$ (5µm, Reprosil) trap column (0.150 mm i.d. x 30 mm) and separated at a flow rate of 1 µL.min$^{-1}$ using a $C_4$ (5 µm, Reprosil) column (0.150 mm i.d. x 400 mm). The following gradient was used: 2% solvent B (20% $H_2O$, 80% ACN, 0.1% FA) from 0–5 min; 20% B at 6 min.; 35% B at 7 min.; 60% B at 16 min.; 99% B from 17–20 min.; and 2% B from 20.2–40 min.

A first LC-MS experiment was acquired at 15,000 resolving power (at m/z 400) with a scan range set to 550-2,000 m/z, 5 microscans (µscans) per MS scan, an automatic gain control (AGC) target value of 5x10$^5$ and maximum injection time of 50 ms. Fragmentation data were recorded using targeted LC-MS/MS experiments. Four precursor charge states were chosen for each protein across their respective charge state distribution and isolated by the quadrupole and subjected to fragmentation with a maximum of two charge states per chromatographic run for each protein (**Table S1**). MS/MS scans were acquired at 120,000 resolving power (at m/z 400) with an isolation width of 1.6 m/z, 5 µscans, an AGC target value of 5x10$^5$ and maximum injection time of 246 ms. Higher-energy collisional dissociation with NCE of 10, 15 and 20% (HCD), electron transfer dissociation with 5, 10 and 15 ms of reaction time and a supplemental higher-energy collisional dissociation with NCE of respectively 5, 10 and 15% (EThcD), ultraviolet photodissociation at 213 nm with 40, 50 and 60 ms of reaction time (UVPD) were used for the fragmentation of intact proteins.

**Table S1.** Targeted m/z and the corresponding charge state for each protein

| Protein | Targeted m/z | Charge state | Protein | Targeted m/z | Charge state |
|---|---|---|---|---|---|
| IGF-I LR3 | 916.965 | 10+ | BCA2 | 763.697 | 38+ |
| | 1013.209 | 9+ | | 829.008 | 35+ |
| | 1139.704 | 8+ | | 906.658 | 32+ |
| | 1302.329 | 7+ | | 1000.357 | 29+ |
| Thioredoxin | 791.973 | 15+ | Protein AG | 935.435 | 54+ |
| | 913.602 | 13+ | | 1010.179 | 50+ |
| | 1079.500 | 11+ | | 1097.943 | 46+ |
| | 1319.240 | 9+ | | 1202.416 | 42+ |
| Protein G | 858.691 | 25+ | Exo Klenow | 756.575 | 90+ |
| | 975.592 | 22+ | | 801.01 | 85+ |
| | 1129.466 | 19+ | | 851.02 | 80+ |
| | 1341.129 | 16+ | | 907.69 | 75+ |

### Data analysis

MS/MS spectral data were first deconvoluted and deisotoped using the Xtract algorithm embedded in FreeStyle$^{TM}$ v1.6.75.20 (Thermo-Scientific) using a fit factor of 80%, a remainder threshold of 25% and a S/N threshold of 3. Scans were averaged across the width of the chromatographic peak. Deconvoluted ion masses were then exported as *.xls files and uploaded into ProSight Lite v1.4 (Fellers *et al.*, 2015) with the appropriate protein sequence. Assigned fragments were finally exported as *.xlsx files. Both *.xls and *.xlsx files were used as input data for TDFragMapper.

### Results

The addition of golden complementary pairs onto the fragmentation map, as displayed in **Figure S1**, is generated instantly when selecting the corresponding option in the filter interface of TDFragMapper. This option makes it easier to the user to localize quickly the golden complementary pairs. A golden complementary pair is a pair of fragment ions (a/x, b/y or c/z) that have been formed by cleavage between the same pair of amino acids (Horn *et al.*, 2000; Kelleher *et al.*, 1999). Since the sum of the masses of the two fragments equals the mass of the targeted protein, golden complementary pairs can greatly enhance confidence in protein characterization (**Figure S2**).

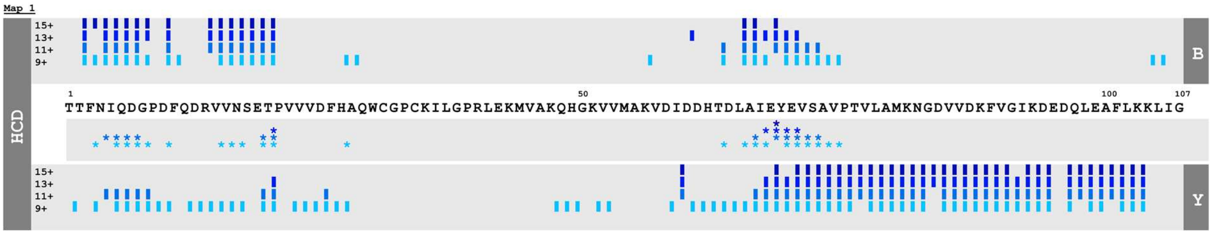


| Precursor charge state | 15+ | 13+ | 11+ | 9+ |
|---|---|---|---|---|
| Residue cleavages (%) | 48 | 51 | 51 | 71 |
| Number of matching fragments | 52 | 59 | 66 | 97 |
| *: Golden complementary pairs | 1 | 5 | 12 | 22 |

**Figure S1: Fragmentation map of human Thioredoxin obtained when varying the precursor charge state in HCD with NCE 20%, with the "Golden complementary pairs" option set on.**

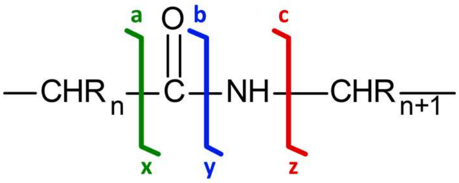


**Figure S2: Golden complementary pairs generated in tandem mass spectrometry of intact proteins**

The overlay of the intensity of deconvoluted fragments with their localization, as displayed in **Figure S3**, is generated instantly by the "Intensity" option embedded in TDFragMapper. This option is of great advantage to quickly spot the most intense fragments for each condition represented on the map. Moreover, variations in the abundance of fragments can be easily spotted. **In Figure S3**, it can be easily seen that the y-fragment between Val74 and Pro75 is the most intense fragment regardless of the selected precursor charge state.

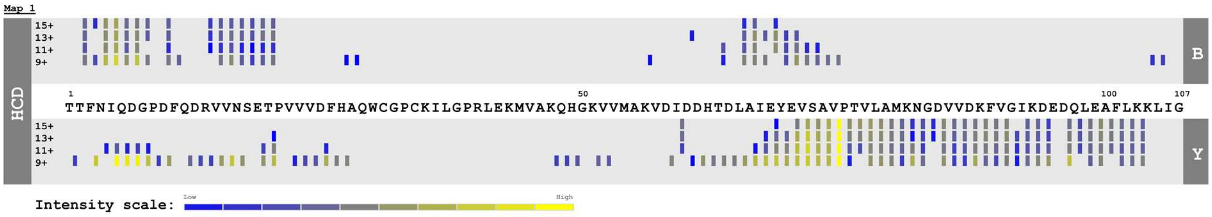


| Precursor charge state | 15+ | 13+ | 11+ | 9+ |
|---|---|---|---|---|
| Residue cleavages (%) | 48 | 51 | 51 | 71 |
| Number of matching fragments | 52 | 59 | 66 | 97 |

**Figure S3: Fragmentation map of human Thioredoxin obtained when varying the precursor charge state in HCD with NCE 20%, with the "Intensity" option set on.**

The complete characterization of proteins by top-down mass spectrometry often requires the use of diverse fragmentation techniques (Fornelli *et al.*, 2018; Brunner *et al.*, 2015). The merging option can be used to sum all the fragments arising from the best fragmentation experiments. A single fragmentation map is created and gathers all a-, b- and c-ions into N-terminal cleavages and all x-, y- and z-ions into C-terminal cleavages. In **Figure S4**, the fragmentation results obtained with HCD (NCE 20%) and with EThcD (5 ms + NCE 5%) are displayed in separate maps. The best fragmentation results, here the fragments obtained with the 9+ precursor in HCD and the 15+ precursor in EThcD, are merged into the final fragmentation map displayed at the bottom with a final residue cleavage of 92%.

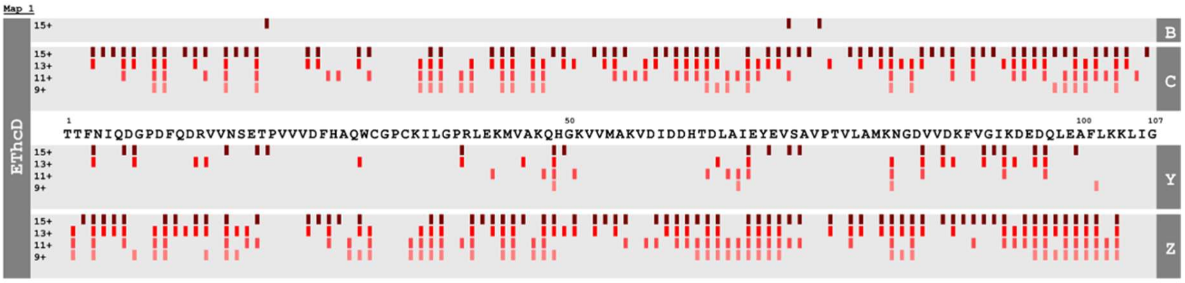


| Precursor charge state | 15+ | 13+ | 11+ | 9+ |
|---|---|---|---|---|
| Residue cleavages (%) | 81 | 70 | 58 | 42 |
| Number of matching fragments | 159 | 133 | 109 | 70 |

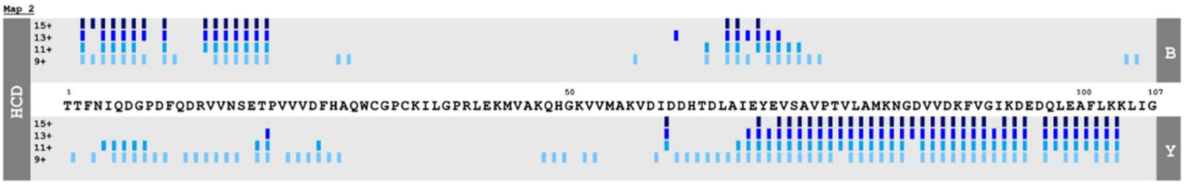


| Precursor charge state | 15+ | 13+ | 11+ | 9+ |
|---|---|---|---|---|
| Residue cleavages (%) | 48 | 51 | 51 | 71 |
| Number of matching fragments | 52 | 59 | 66 | 97 |

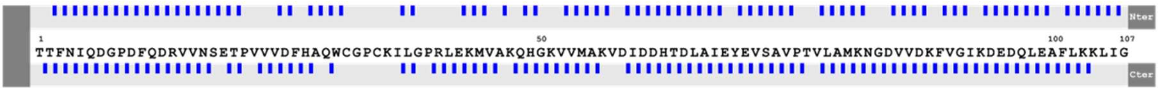


Residue cleavages: 92%

**Figure S4: Fragmentation maps of human Thioredoxin obtained when varying the precursor charge state in EThcD with 5ms + NCE 5% (top), in HCD with NCE 20% (middle) and with the best results of EThcD and HCD combined using the "Merging" option (bottom).**
